# SUB-OPTIMAL ENVIRONMENTAL CONDITIONS PROLONG PHAGE EPIDEMICS IN BACTERIAL POPULATIONS

**DOI:** 10.1101/2022.10.28.514181

**Authors:** Henry Goehlich, Olivia Roth, Michael Sieber, Cynthia M. Chibani, Anja Poehlein, Jelena Rajkov, Heiko Liesegang, Carolin C. Wendling

## Abstract

Infections by filamentous phages influence bacterial fitness in various ways. While phage-encoded accessory genes, e.g., virulence genes, can be highly beneficial, the production of viral particles is energetically costly and often reduces bacterial growth. Consequently, if costs outweigh benefits, bacteria evolve resistance which can shorten phage epidemics. Abiotic conditions are known to influence the net-fitness effect for infected bacteria. Their impact on the dynamics and trajectories of host resistance evolution, however, remains yet unknown. To address this, we experimentally evolved the bacterium *Vibrio alginolyticus* in the presence of a filamentous phage at three different salinity levels, i.e., (1) ambient (2) 50% reduction, and (3) fluctuations between reduced and ambient. In all three salinities, bacteria rapidly acquired resistance through super infection exclusion (SIE), whereby phage-infected cells acquired immunity at the cost of reduced growth. Over time, SIE was gradually replaced by evolutionary fitter surface receptor mutants (SRM). This replacement was significantly faster at ambient and fluctuating conditions compared to the low saline environment. Our experimentally parameterized mathematical model explains that suboptimal environmental conditions, in which bacterial growth is slower, slow down phage resistance evolution ultimately prolonging phage epidemics. Our results imply that, if filamentous phages encode virulence genes, these may persist longer in bacterial populations at sub-optimal environmental conditions, which, in times of climate change, are becoming more frequent. Thus, our future ocean may favour the emergence of phage-born pathogenic bacteria, and impose a greater risk for disease outbreaks, impacting not only marine animals but also humans.

## INTRODUCTION

Parasites and pathogens, which often impose a strong fitness burden on their host, are the strongest evolutionary force across the living world (Ebert & Hamilton, 1996; Keesing et al., 2010; Thompson, 1998). While the evolution of resistance is a common strategy for hosts to counteract parasitism, the costs of resistance often vary, making it difficult to predict when and how resistance evolves. Theory predicts and experiments confirmed that in the absence of significant resistance costs, resistance rapidly emerges and reaches fixation (Agrawal & Lively, 2002; Buckling & Rainey, 2002; Wendling et al., 2022). With increasing costs of resistance however, intermediate resistance levels or co-existence of non-resistant and resistant subpopulations may appear (Boots & Haraguchi, 1999).

The surrounding abiotic environment adds complexity to the interaction between hosts and parasites. That is, because resistance costs often depend on environmental factors leading to different evolutionary outcomes *in vivo* compared to *in vitro* experiments (Hernandez & Koskella, 2019). In addition, variations in the abiotic environment not only affect resistance costs directly but impact the fitness of the interacting species to various degrees, which further complicates evolutionary predictions; for a review see (Wolinska & King, 2009). We predict that environmental conditions that increase parasite burden should select for faster resistance evolution. On the other hand, environmental conditions that decrease host fitness, for instance by reducing host generation times, might decelerate or even constrain resistance evolution and could thereby prolong epidemics.

To test these predictions, we studied how changing environmental conditions influence the trajectory of resistance evolution. To do so, we performed a selection experiment at three different salinity levels using the marine bacterium *V. alginolyticus* strain K01M1 (Chibani, Roth, Liesegang, & Wendling, 2020) and the filamentous *Vibrio* phage VALGΦ8 (Chibani, Hertel, Hoppert, Liesegang, & Wendling, 2020). Filamentous phages (family *Inoviridae*), i.e., long, thin proteinaceous filaments which contain a circular single-stranded DNA genome, are ideal model systems to study virulence (Messenger, Molineux, & Bull, 1999; Turner, Cooper, & Lenski, 1998) and resistance evolution (Wendling et al., 2022). Filamentous phages often establish chronic infections whereby virions are continuously released without lysis. This costly feature of filamentous phages often reduces bacterial growth rates (Wendling et al., 2022), because bacteria pay the metabolic costs resulting from phage replication and through phage-encoded proteins inserted into the bacterial membrane (Mai-Prochnow et al., 2015).

In a previous study (Wendling et al., 2022) we found that in the presence of *Vibrio* phage VALGΦ8 the bacterium *V. alginolyticus* K01M1 evolved phage resistance by modifying the viral binding site. These mutants rapidly replaced phage-carriers, which were resistant due to super infection exclusion (SIE)-immunity. This replacement happened because intracellular phage presence, which reduced bacterial growth, was more costly than surface receptor mutations (SRM). However, whether filamentous phages are costly (Bull, Molineux, & Rice, 1991; Shapiro & Turner, 2018; Shapiro, Williams, & Turner, 2016) or beneficial (Hay & Lithgow, 2019; Ilyina, 2015) largely depends on the environmental context.

Environmental variations impact marine microbes in manifold ways (for a review see (Cavicchioli et al., 2019). One parameter that is constantly fluctuating, in particular in coastal waters due to changes in rainfall, is salinity. Bacteria living in such fluctuating salinity regimes, such as *Vibrionaceae* have thus developed various ways to cope with osmotic stress (Gregory & Boyd, 2021). However, sub-optimal salinity levels, characterized by reduced bacterial growth, can increase bacterial susceptibility to infections with filamentous phages (Goehlich, Roth, & Wendling, 2019). This suggests that changing salinity levels will change the trajectory and tempo of phage resistance evolution and phage epidemics, i.e., phage presence, in bacterial populations. Such insights will allow us to make better predictions of bacterial epidemics, as several infectious diseases are caused by virulence genes encoded on filamentous phages. Upon integration into the bacterial genome these genes can turn bacteria into deadly disease agents (Faruque & Mekalanos, 2012).

Combining experimental evolution and mathematical modelling across different levels of salinity, we found that SIE persisted longer, and resistant mutants emerged later at sub-optimal salinity levels. Our experimentally parameterised mathematical model suggests that salinity-dependent bacterial growth is the driving factor behind these different trajectories: optimal growth conditions select for fast resistance evolution and rapid replacement of SIE by SRM, whereas sub-optimal conditions constrained resistance evolution and allowed filamentous phages to persist longer in bacterial populations. Taken together, our data suggest, that in the case of filamentous phages that encode virulence genes, sub-optimal environmental conditions, which will become more frequent with climate change, may favour the emergence of phage-born pathogenic bacteria, and impose a greater risk for disease outbreaks in eukaryotes.

## MATERIAL AND METHODS

### (a) Study organisms

*Vibrio alginolyticus* strain K01M1 (Gene bank accession number: CP017889.1, CP017890.1) and *V. alginolyticus* strain K04M1 harbouring the focal filamentous *Vibrio* phage VALGΦ8 (MN690600) were isolated from healthy pipefish (Roth, Keller, Landis, Salzburger, & Reusch, 2012) and characterised by whole genome sequencing (Chibani, Roth, et al., 2020). The *V. alginolyticus* strain K01M1 harbours a resident filamentous *Vibrio* phage VALGΦ6 which replicates at a lower frequency in comparison to *Vibrio* phage VALGΦ8 (Chibani, Hertel, et al., 2020).

### (b) Selection experiment

We obtained *Vibrio* phage VALGΦ8 from overnight cultures (ONCs) of K04M1. ONCs were centrifuged for 5 min at 6000g. Phage-containing supernatant was filtered (pore size: 0.20 μm) to remove all bacteria, 10-fold diluted in TM buffer (modified from Sen & Gosh (2005): 50% (v/v) 20 mM MgCl2, 50% (v/v) 50 mM Tris–HCl, pH 7.5) and subsequently stored at 4° C overnight.

In parallel, we initiated six replicate populations from six independent clones of *V. alginolyticus* K01M1 for each treatment. The treatments comprised a set of 18 populations co-evolving with the filamentous *Vibrio* phage VALGΦ8 at three different salinity conditions: a) constant low (7 Practical Salinity Unit (PSU)), b) constant ambient (15 PSU), and c) fluctuating (starting with 7 PSU on transfer 0 and 1 and then alternating between 7 and 15 PSU every transfer). Another set of 6 populations were propagated at each same salinity level but without added phages as controls.

Populations were established from 60 µl overnight cultures (∼5×10^9^ CFU/ml) and inoculated with 300 µl *Vibrio* phage VALGΦ8 suspension (∼5×10^8^ phage particles/ml) in microcosms (30 ml glass container with loose plastic lid) containing 6 ml of liquid medium (medium101: 0.5% (w/v) peptone, 0.3% (w/v) meat extract, 0.7% (w/v) or 1.5% (w/v) NaCl in MilliQ water). Microcosms were incubated at 25 °C and 180 rpm. Every 24 hours, 60 µl of each population was transferred into fresh microcosms with 6 ml medium to allow continuous growth. Each transfer is equivalent to approximately 7.5 generations, presuming that all populations diluted 1:100 reach stationary phase after 24 hours, which we confirmed (Supplementary material, Figure S1).

Initially, phage and bacterial densities were determined on transfer T0, T1, T2, T3, T4. Subsequently, we quantified phage and bacterial densities using a 1-1-0 rhythm, i.e., two consecutive transfers, followed by a transfer with no density measurements. Using this rhythm, in contrast to every 2^nd^ day sampling, allowed us to include both salinity levels of the fluctuating treatment in our measurements. On transfer T0, T2, T3, T4, T6, T10, T12, T16, T22, and T29 whole bacterial population samples, phage populations and 24 randomly isolated single colonies from each bacterial population were frozen at -80° C at a final concentration of 33% glycerol. Two populations from the control treatment that tested positive for VALGΦ8 infection, indicating contamination, were excluded from later assays.

### (c) Bacterial and phage densities

#### Bacterial densities

We determined bacterial densities by plating 100 µl of a dilution of 10^−6^ or 10^−7^ depending on optical density of the culture on *Vibrio* selective Thiosulfat-Citrat-Bile-Saccharose (TCBS; Fluka Analytica) agar plates. Plates were incubated over night at 25° C and the total number of colonies was counted the following day.

#### Phage densities

We used spectrophotometry to quantify the prevalence of the filamentous phages following a protocol published on the website of Antibody Design Labs™ (Abdesinglabs, 2013) with some modifications (Wendling 2022). This method, which is based on small-scale precipitation of phages by single PEG precipitation is the standard method for quantifying filamentous phages because infection by filamentous phages does not necessarily result in lysis of bacterial cells making it impossible to quantify plaque forming units per ml using standard spot assays. The quantification limit for this method is 1.3^12^ phage ml^-1^, which was calculated by adding ten times the standard deviation to the average of the blanks (Shrivastava & Gupta, 2011). Below this threshold, the spectrophotometric quantification of VALGΦ8 is susceptible to inaccuracies.

In parallel, we used standard spot assays to determine the presence or absence of phages as PEG values were often below the quantification limit. Here, we observed opaque zones of reduced bacterial growth, which are, however, not necessarily caused by lysis and are mainly due to reduced growth rates. To do so, we diluted all populations at the end of each transfer 1:100 using medium101 with the respective salinity and followed a protocol for spot assays that we developed during a previous study (Wendling et al., 2017).

### (d) Overnight growth curves of bacterial populations

For transfer 1, we additionally monitored the optical density of a subculture of each evolving population to confirm, that despite a lower growth rate at 7 PSU, populations still reached stationary phase. Approximately 1 h after the daily transfer of 60 µl subpopulation into the fresh media, we added a 200 µl subpopulation of the new culture to a transparent, flat-bottom 96-well microtiter (Nunclon™) plate and generated 23h growth curves in a TECAN infinite M200 plate reader, where optical density measurements were taken every 20 min at 600 nm (OD_600_).

### (e) Measuring phage defence

We quantified bacteria that could not get infected by the ancestral *Vibrio* phage VALGΦ8 by measuring the phage’s ability to inhibit bacterial growth using a reduction in bacterial growth assay (RBG) in liquid culture adapted from (Poullain, Gandon, Brockhurst, Buckling, & Hochberg, 2008) and modified according to (Wendling, Goehlich, & Roth, 2018). To do so, we took 24 random colonies of each infected population and 12 random colonies of each control population from transfers T0, T2, T3, T4, T6, T10, T12, T16, T22 and T29. In a few cases the number of tested clones was below 24.

### (f) Frequency of phage-carrying strains

In a previous study (Wendling et al., 2022), we found two alternative defence strategies conferring resistance against the co-evolving filamentous *Vibrio* phage VALGΦ8: (1) superinfection exclusion (SIE) resulting from the presence of the co-evolving phage in the bacteria cell or (2) surface receptor mutations (SRM) in the MSHA type IV pilus preventing phage attachment. To quantify the frequency of co-evolving strains that acquired resistance by superinfection exclusion, we used colony PCR with a primer pair that specifically targets a unique region in the co-evolving *Vibrio* phage VALGΦ8 following a previous protocol (Wendling et al., 2022). PCR-bands at a length of 1.2 kbp were interpreted as presence of VALGΦ8 in the bacterial cell indicating SIE. The absence of a PCR product in resistant clones was interpreted and later confirmed by whole genome sequencing as SRM.

Based on these PCR results, we selected one PCR-positive and one PCR-negative clone of every population from transfer six as well as one clone from each control population. We chose transfer six because it was the last time point of the experiment at which at least one of the 24 picked clones of the co-evolving population was either PCR positive or PCR negative (with one exception). Selected clones were then used for (a) phenotypic assays including bacterial growth rate and phage production and (b) whole genome sequencing.

### (g) Phenotypic assays

#### Growth rates of single clones

Absolute Growth rate of single clones: Overnight cultures were diluted 1:100 and growth rate was measured by generating 24 h growth curves using an automated plate reader (TECAN infinite M200).

#### Phage production of single clones

Overnight cultures were diluted 1:100 and incubated in 15 ml Falcon tubes containing 6 ml medium101 and the salinity the clones were exposed to during the selection experiment. Clones evolved in the fluctuating salinity treatment were incubated at 15 and 7 PSU. After 24 h at 25° C and 180 rpm, phage density was measured using spectrophotometry following PEG precipitation as described above.

### (h) Whole genome sequencing of evolved clones

To determine the genetic basis of the observed resistance mechanism, we sequenced the genomes of the same clones (2.7.), which were used for the fitness assays, i.e., one phage-resistant PCR-negative clone and one phage-resistant PCR-positive clone per population at transfer six. We also included one phage-susceptible clone from each control population. DNA for sequencing was isolated from cultures grown in medium101 for 16 h at 25° C and 180 rpm. High molecular weight DNA was extracted using the DNeasy Blood & Tissue Kit (Qiagen, Hilden, Germany).

Sequencing was done using an Illumina 2500 platform. The quality of the reads was evaluated with the program FastQC Version 0.11.5. Illumina reads of each sampled clone were assembled using Spades version 3.14.0 (Bankevich et al., 2012). In-silico PCR (http://insilico.ehu.eus) on assembled contigs as template and VALGΦ8 phage-specific primers were used to confirm the empirical PCR results and the presence of the infecting phage in each clone. To identify mutations conferring resistance against the VALGΦ8 we used the *breseq* Variant Report pipeline (version 0.35.0) (Deatherage & Barrick, 2014). Paired reads were mapped to the annotated reference genome of the ancestral K01M1 strain (GenBank accession number: CP017889.1, CP017890.1). Variants that occurred in all replicates were discarded as they represent differences between the reference genome and the ancestral genotype used in the experiment.

### (i) Statistical analyses

All statistics were performed in the R 3.5.1 statistical environment (RCoreTeam, 2022). We excluded the contaminated control populations P3 and P15 from all further analyses. If not otherwise stated, we used Maximum Likelihood estimation in linear mixed effect models and tested significance using ANOVA type marginal for models with interaction, or ANOVA type sequential in models with no interaction. We initially included all interaction terms in the models before reducing each model by sequentially removing non-significant interactions according to Akaike Information Criterion (AIC) (Akaike, 1974). Post-hoc tests were carried out using TukeyHSD (package: multcomp) or estimated marginal means (package: emmeans) for generalized least square models. We report mean plus standard error in the results section and the figures unless otherwise stated.

#### Bacteria and phage dynamics

We analysed bacteria and phage dynamics over time using generalised least square (gls) models (package: nlme4). The models were fitted using *Phage* (VALGΦ8 presence/absence) and *Salinity* (7 PSU, 15 PSU, or fluctuating) as fixed effects, and *Transfer* as integer. Weight *varIdent(form=∼1*|*Transfer)* was introduced to account for heterogeneous variances over time and the term (*corAR1(0*.*6, form=∼1*|*Day*) for autocorrelation. The model suggests a significant three-way interaction between *Phage, Transfer* and *Salinity* for bacterial dynamics (gls: *F*_1,665_ = 5.2, *p* = .006; Supplementary material, Table S1) and a significant interaction between *Phage* and *Transfer* for phage dynamics (gls: *F*_1,678_ = 3.3, *p* < .039; Supplementary material, Table S2). Thus, we excluded control populations (without phages) to test for the effect of *Salinity* and *Transfer* on bacteria and phage dynamics. To compare the decrease in phage particles between the different salinities from the time point of the highest phage concentration to the time point with phage concentrations below quantification level, we run an additional gls model comprising the transfers T3 to T9 only.

#### Reduction in bacterial growth assay

Bacterial resistance, i.e., the ratio of susceptible to resistant clones was analysed using a generalized linear mixed model (glmer – package lme4) with a binomial error distribution, *Salinity* (7 PSU, 15 PSU, fluctuating), *Phage* (VALGΦ8 presence/absence) as fixed effects, *Transfer* as integer and *Population* as random effect. The model suggests a significant interaction between *Phage* and *Transfer* (glmer: χ^2^ _2_ = 9.8, p = .004, Supplementary material, Table S3). We thus excluded control populations (without phages) from further analyses to test for the effect of *Salinity* and *Transfer* on phage-resistance evolution.

#### Resistance mechanism

We analysed the proportion of resistant clones carrying VALGΦ8 in the evolving populations as a ratio of resistant phage-infected (PCR positive) vs. resistant non-infected (PCR negative) clones using a generalized linear mixed model (glmer) – package lme4 with a binomial error distribution. *Salinity* (7 PSU, 15 PSU, fluctuating) and *Transfer* were assigned as fixed effects and *Population* as random effect.

#### Fitness effects

To determine differences in the number of free phages produced per culture and in growth rates between ancestral strains and evolved strains and between both resistance forms, we used Welch’s pairwise t-tests with sequential Bonferroni corrections. We further performed a Spearman’s rank correlation analysis to determine whether phage production impacted bacterial growth rates.

### (j) Mathematical model of resistance evolution

We employed a special case of the mathematical model presented in Wendling et al. 2022. In short, the model describes the dynamics of the ancestral clones (with density *B*), resistant SIE clones (*I*), resistant SRM clones (*R*), SIE clones that have also acquired the MSHA mutation (*IR*), and the phage population (*V*) in batch cultures. The SIE clones (I and IR) have a reduced growth rate relative to the ancestor due to a fixed virulence factor caused by intra-cellular production of the virus. The fitness effects of different salinities are reflected by different ancestral growth rates. To keep the model as simple as possible we did not model subpopulation dynamics under fluctuating conditions. The model equations and parameters can be found in the Supplementary Information and Table S4.

## RESULTS

### Ecological dynamics vary across salinities

We experimentally evolved six replicate populations of the bacterium *Vibrio alginolyticus* K01M1 with or without the filamentous *Vibrio* phage VALGΦ8 for 30 serial transfers (∼ 240 bacterial generations) at three different salinity treatments: ambient (15 PSU), low (7 PSU) and periodically fluctuating between 15 and 7 PSU every 24 hours.

In the absence of phages, bacterial densities differed significantly between salinity treatments (significant Salinity:Transfer interaction in gls: F_1,313_ = 3.7, *p* = 0.026; Supplementary material, Table S5, Figure S2). When exposed to 15 PSU or fluctuating salinities population densities remained constant at around 6.3×10^9^ CFU/ml ± 2.1×10^8^ (mean ± s.e. in 15 PSU) and 5.4×10^9^ CFU/ml ± 2.0×10^8^ (fluctuating treatment) throughout the experiment (no significant Transfer effect in gls at 15 PSU: F_1,117_ = 0.3, *p* = 0.574; and fluctuating salinities: F_1,101_ = 1.9, *p* = 0.164). However, at 7 PSU bacterial densities were significantly lower at the beginning of the experiment i.e., transfer 1 (T1) - T3 (3.0×10^9^ ± 4.1×10^8^) compared to the end i.e., T25 - T29, where they reached levels comparable to the other two salinity treatments (6.3×10^9^ CFU/ml ± 3.7×10^9^; significant Transfer effect in gls at 7 PSU: F_1,95_ = 10.0, *p* = 0.002). This confirms that low salinities are stressful, reducing bacterial growth rates but also suggests that bacteria found a way to cope with this stress.

Similarly, bacterial densities varied significantly in the presence of phages across the three different salinity treatments (significant Salinity:Transfer effect in gls: *F*_2,346_ = 3.6, *p* = .030; Figure 1a; Supplementary material, Table S6). While phages reduced bacterial densities in all salinity treatments by several orders of magnitude, we observed substantial differences between treatments regarding (i) time point of lowest bacterial densities, (ii) lowest total population size, and (iii) time point of recovery (i.e., population size is comparable to T0). At 15 PSU bacterial densities reached a minimum 48 hours post infection [hpi] (6.9×10^8^ CFU/ml ± 8.8×10^7^), at fluctuating salinities 72 hpi (4.4×10^8^ ± 6.3×10^7^ CFU/ml) and at 7 PSU 96 hpi (1.45×10^8^± 4.2×10^7^).

**Figure 1:**
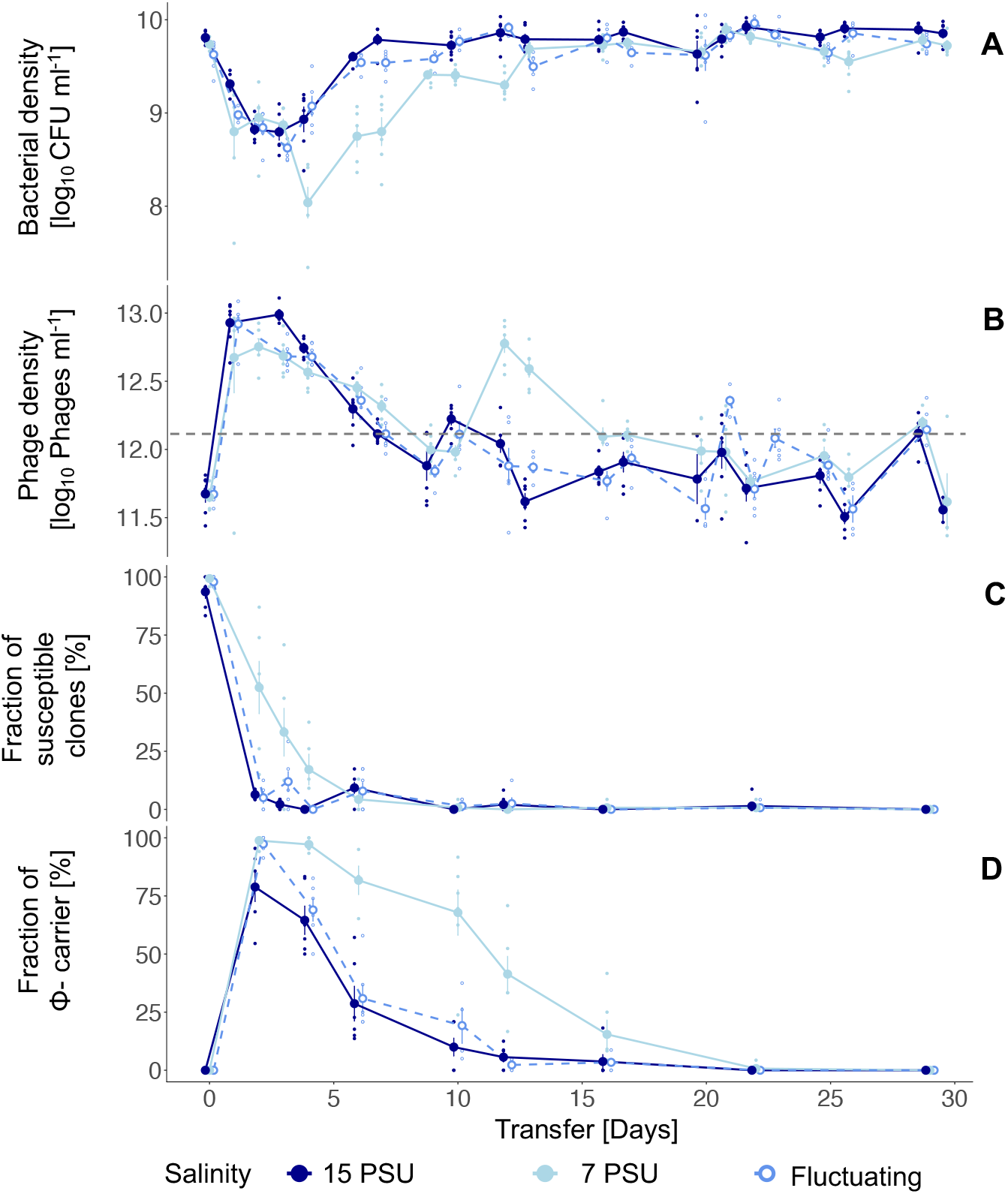
Population dynamics in the presence of VALGΦ8 over 30 transfers. **(A)** Bacterial densities are shown as colony forming units (log_10_ CFU ml^-1^) and **(B)** Phage densities as phage particles measured by PEG precipitation (log_10_ phage particles ml^-1^). The black dashed horizontal line in panel B indicates the quantification limit for phage concentrations. **(C)** Fraction of phage susceptible clones (n=24): Susceptibility was tested against the ancestral phage VALGΦ8. **(D)** Fraction of Φ-carrier within resistant clones. In all panels, individual replicate populations are represented by small dots (n=6) and means by larger points (± s.e.). Colours represent the salinity level during the evolution experiment (dark blue: 15 PSU, light blue: 7 PSU, cobalt blue and dashed line: fluctuating).

Correspondingly, phages amplified massively and rapidly in all salinity treatments reaching a maximum 24 hpi (Figure 1b). This immediate peak was higher in 15 PSU (8.96×10^12^ ± 1.2×10^12^ phage particles per ml [PP/ml]) and fluctuating salinities (8.71 × 10^12^ ± 1.1×10^12^ PP/ml) compared to 7 PSU (7.2 × 10^12^ ± 1.4×10^12^ at 7 PSU) 24 hpi (Figure 1b). This suggests that the initial decline in bacterial population densities (Figure 1a) directly resulted from massive phage particle production, which is costly for the host cell. Note, quantified phage particles in control populations resulted from the resident filamentous *Vibrio* phage VALGΦ6 that produces particles at a low background rate which mainly remained below the quantification limit throughout the experiment.

Over time, phage exposed bacterial populations recovered to starting densities around T6/T7 at 15 PSU and fluctuating salinities. However, at 7 PSU, time to recovery took twice as long (T12). Bacterial recovery was accompanied by a gradual decrease in phage densities in all salinities, but this decline was slowest at 7 PSU (significant Salinity:Transfer effect in gls: *F*_2,348_ = 6.0, *p* = .004; Supplementary material, Table S7).

### Bacterial resistance acquisition is delayed at 7 PSU

These opposite ecological dynamics of bacteria and phage populations suggest that bacteria acquired a defence mechanism that enabled the recovery of their populations. Indeed, susceptible clones rapidly declined and did so significantly faster in 15 PSU and fluctuating salinities (significant *Transfer:Salinity* effect in generalised linear mixed effect model (glmer): χ^2^ _2_ = 9.7, *p* = .008; Supplementary material, Table S8; Figure 1c; Tukey’s HSD, 15 PSU – 7 PSU: *p* < 0.001; Fluctuating – 7 PSU: *p* = 0.002, Supplementary material, Table S8a), where resistance reached fixation at T4 (Figure 1c). In contrast at 7 PSU, fixation of resistance took significantly longer and reached fixation in five out of six populations at T10 and in all populations at T29.

K01M1 can acquire resistance against VALGΦ8 via two distinct mechanisms (1) super infection exclusion (SIE) immunity, where phage-infected cells are protected from subsequent infections at the cost of reduced bacterial growth or (2) surface receptor mutations (SRM) at the MSHA-pilus which serves as an entry receptor for VALGΦ8 at the cost of reduced motility (Wendling et al., 2022). To determine the frequency of SIE and SRM we used PCR with specific primers that target VALGΦ8. We found that SIE rapidly increased in frequency and dominated the populations in 15 PSU and fluctuating salinities after 48 hpi and in 7 PSU after 96 hpi. This rapid emergence was followed by a gradual decline of SIE carriers, which was significantly faster in 15 PSU and in fluctuating salinities compared to 7 PSU (significant *Salinity:Transfer* interaction in generalised linear mixed effect model (glmer): χ^2^ _2_ = 32.0, *p* < .001; Supplementary material, Table S9 and Tukey’s HSD, 15 PSU – 7 PSU: *p* < .001; Fluctuating – 7 PSU: *p* < .001; Supplementary material, Table S10; Figure 1d). Since all populations contained almost no susceptible bacteria 48 hpi in 15 PSU and fluctuating salinities, and 96 hpi in 7 PSU, the observed decline in Φ-carriers suggests their replacement by emerging SRM, which was more strongly selected for at 15 PSU and in fluctuating salinities, i.e., 90% SRM at T12 compared to T22 in 7 PSU.

### Resistance is associated with mutations in MSHA type IV pilus encoding genes

To confirm that the displacement of SIE was driven by emerging SRM, as previously described in (Wendling et al., 2022), we randomly chose one PCR-positive and one PCR-negative clone from each population for whole genome sequencing at T6. We first performed in silico PCRs with virtual primers targeting VALGΦ8 on all sequenced clones and found that three clones which were initially assigned as PCR-negative did carry VALGΦ8, whilst one clone that was initially classified as PCR-positive did not carry VALGΦ8. Subsequent data analysis was done using the classification based on the in-silico PCR. In addition, we found a third genotype: DNA-samples from five PCR-positive clones from the fluctuating treatment contained VALGΦ8 DNA but at concentrations lower than the genomic DNA (Supplementary material, Figure S3). Thus, we assume that these five clones are associated with non-infecting phages and that the low concentration of VALGΦ8-DNA reflects a dilution of phages particles over evolutionary time. In addition, these five genomes carried mutations in the *mshA*-operon that have previously been associated with phage resistance (Wendling et al., 2022). These clones are from here on referred to as Φ*-particle associated-SRM (*Φ*_SRM*).

After filtering out mutations that occurred in control treatments, we found no loci with mutations on chromosome 2, the plasmid pl9064, the resident phage VALGΦ6, or the coinfecting phage VALGΦ8. However, on chromosome 1, we identified 13 loci with mutations that were absent in clones from the control treatment suggesting an association with phage-mediated selection (Supplementary material, Table S11). Of these 13 loci, six loci were exclusive to SIE clones but randomly distributed across salinities. These included several SNPs and insertions causing frame-shift mutations in three different intergenic regions of which the sensor histidine kinase RcsC / hypothetical protein (K01M1_16000) has been repeatedly hit but whose function we cannot explain.

The remaining three loci exclusive to SIE clones comprised two hypothetical proteins and the bvgS protein, each of which has been hit only once. Three of the 13 loci were exclusive to PCR negative clones and Φ_SRM. These included a twitching motility protein (*pilT*) and two genes belonging to the MSHA type IV pilus operon (*K01M1_28150, K01M1_28240, sct_C*, Supplementary material, Table S11). Indeed, all but one, PCR negative clones acquired substitutions, duplications, insertions, or deletions in one of five different genes belonging to the MSHA type IV operon (*K01M1_28140, K01M1_28150*, and *K01M1_28240, epsE_2*, and *sct_C*, Figure 2 and Supplementary material, Table S11). Among these seven SNVs caused severe frame-shift mutations which presumably have a high impact of the functioning of these proteins. This is in line with our previous findings (Wendling et al., 2022) suggesting strong parallel resistance evolution against the filamentous phage VALGΦ8 resulting from modifications of the phage entry receptor, i.e., the MSHA type IV pilus.

**Figure 2:**
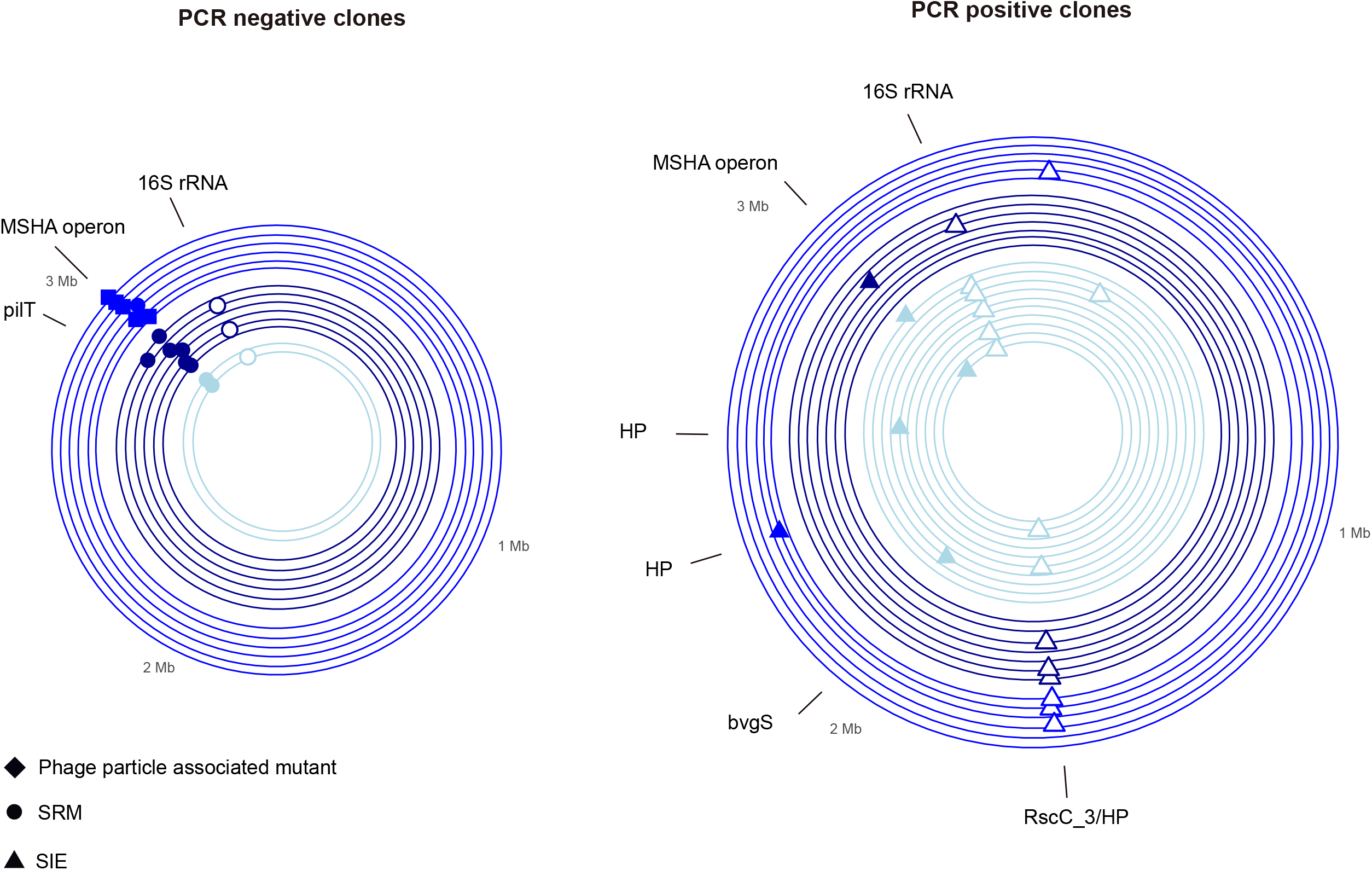
Genetic loci in chromosome 1 under positive selection as indicated by parallel genomic evolution in populations exposed to phages and different salinity regimes (light blue: 7PSU, dark blue: 15 PSU, cobalt blue: fluctuating). *Right*: PCR positive clones, i.e., SIE clones (triangles). Left: PCR negative clones, i.e., SRM (circles) or Φ_SRM (diamonds). Only loci which are not present in control populations are shown. Concentric circles correspond to one clone isolated from each replicate population per salinity. Mutation events in coding regions are shown by filled shapes, mutation events in non-coding regions by open shapes. For more details on the underlying mutations see (Supplementary material, Table S11). HP = Hypothetical Protein

Note, filtering for salinity-specific results did not reveal any parallel patterns that could explain a genetic adaptation to the 7PSU treatment.

### Net bacterial growth rate determines the rate of resistance evolution in a mathematical model

To generalize our findings across more than two salinity levels and to identify the driver behind the delayed resistance evolution at 7 PSU we modified an existing parameterized, mathematical model (Wendling et al, 2022). As in the selection experiment, bacterial population densities declined upon phage infection (Supplementary material, Figure S4a). By simulating phage selection across different salinity levels, we found that this decline was more pronounced and prolonged at 7 compared to 15 PSU (Supplementary material, Figure S4a). Bacterial populations recovered after resistant sub-populations, initially comprised almost exclusively of SIE clones, quickly became dominant (Supplementary material, Figure 4c). As in our experimental data, the model shows that SRM eventually replace SIE clones and become the dominant resistance mechanism (Supplementary material, Figure 4d).

The replacement of SIE clones by SRM is due to the SRM clones’ higher growth rate irrespective of the salinity the population was evolving at (t-test 15 PSU: *t* _6.7_ = 3.14, *p* = 0.02; t-test 7 PSU: t_9.8_ = 6.5, *p* <0.001; Figure 3a). This observed lower growth rate of SIE clones is driven by higher levels of phage production, which was significantly higher for SIE compared to SRM in 15 PSU (t-test: *t* _5.3_ = -2.5, *p* = 0.05), however not in 7PSU (t-test: *t* _4.01_= -0.88, *p* = 0.43; Figure 3b). In both salinities, increasing phage production significantly impaired bacterial growth which was stronger at 15 PSU (rho= -0.67, p=0.017) compared to 7 PSU (rho=-0.63, p=0.032, Figure 3c).

**Figure 3:**
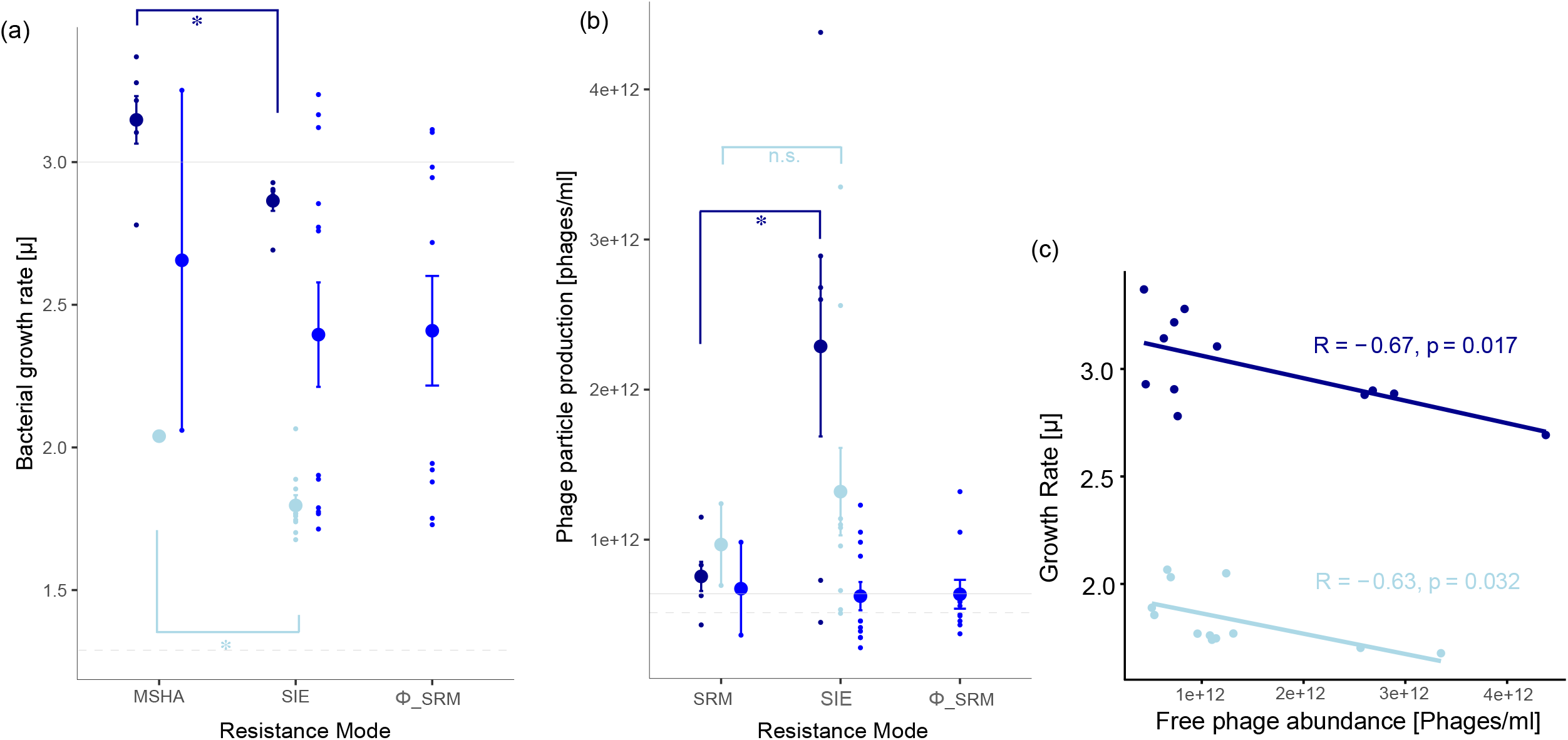
(a) Growth rate µ and (b) Phage particle production [phages/ml] for the three different resistance types: SIE, SRM, and Φ-SRM. Solid and dashed horizontal lines correspond to the ancestral K01M1 strain grown at 15 and 7 PSU respectively. Note: Phages from the ancestral K01M1, from SRM, and the Φ-SRM clones stem from an ancestral filamentous *Vibrio* phage VALGΦ6 integrated on chromosome 2 of K01M1 (Chibani, Roth, et al., 2020). Shown are means ± s.e., n=24. (c) Correlation between bacterial growth rate [µ] and production of free phages measured as phage particles per ml per clone for 7 and 15 PSU.

Consistent with the experimental data, our model predicts a faster replacement of SIE by SRM in 15 than in 7 PSU (Supplementary material, Figure 4d). This is due to the higher growth rate in 15 PSU and increased selection for the combination of fast growth and phage-resistance characteristic for SRM clones compared to SIE clones. This effect holds across a range of growth rates, with lower growth rates generally delaying fixation of SRM clones (Figure 4). This delay at lower growth rates is more pronounced when the phage production rate is low, as this results in lower phage densities and reduced selection for phage resistance (Figure 4).

**Figure 4:**
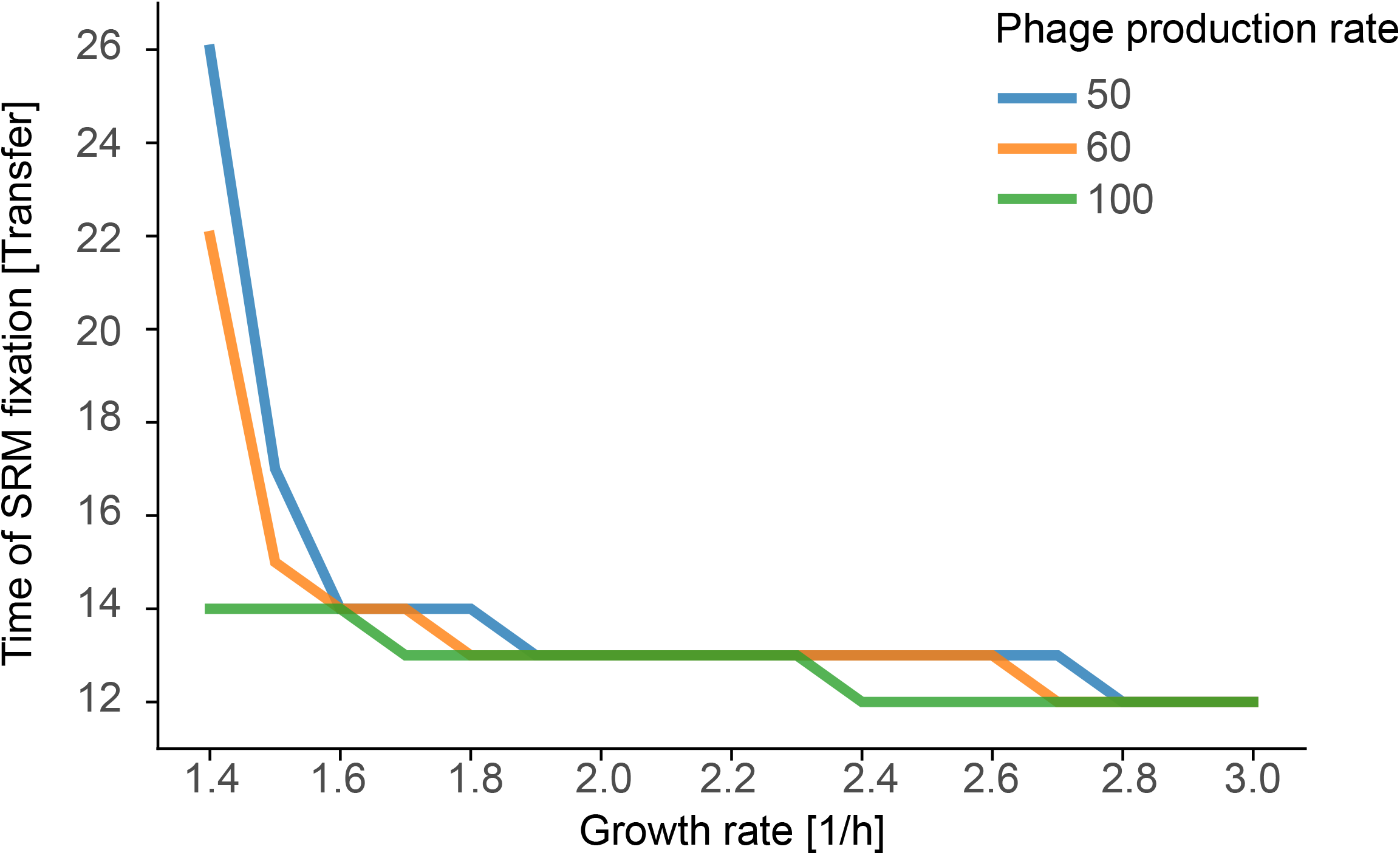
Time of fixation of SRM clones, defined as the transfer at which SRM clones first make up more than 99% of the total population, as a function of growth rate. Higher growth rates speed up the spread of SRM clones, indicated by dominance of SRM clones at earlier transfers. This effect is more pronounced for lower phage production rates.

## DISCUSSION

Changing environmental conditions can have profound implications on species interactions and their co-evolutionary dynamics (Northfield & Ives, 2013). Exposing bacteria to a filamentous phage across different salinity regimes, we found that SIE immunity appeared rapidly and at a similar rate irrespective of salinity. Over time, SRM replaced SIE, due to a selective advantage. However, selection for SRM over SIE was slower in 7 compared to 15 PSU. This is, because at 15 PSU, bacteria have a higher growth rate allowing phages of SIE clones to produce more phage particles which then resulted in a higher fitness cost of SIE clones at optimal compared to suboptimal or fluctuating salinity conditions.

To simulate the relationship between salinity dependent growth and how this influences phage production and its associated costs during selection for resistance we included a mathematical model in which we varied growth rate and phage production across a wider range. The model confirmed that increasing growth rates, which one can expect at optimal environmental conditions, result in higher phage production and thus a faster replacement of SIE by SRM. Thus, as in the experiment, sub-optimal environmental conditions allowed SIE to persist longer in the population which ultimately prolonged the epidemic.

In the costal ecosystems of the Kiel Fjord, where our model organisms originate from, salinity fluctuations due to rainfall, river run offs, or water inflow from the North Sea, by up to 6 PSU throughout the year or occasionally within a few days are common (Bock & Lieberum, 2017; Girjatowicz & Swiatek, 2016). In addition, climate change scenarios predict a salinity drop by up to 5 PSU for this region until the end of the century (Meier, Kjellstrom, & Graham, 2006). Marine bacteria, which have evolved to grow optimally at high salinities, can cope with osmotic changes using a range of different mechanisms to maintain cellular homeostasis; for a review see (Gregory & Boyd, 2021). Given this pre-adaptation to such fluctuations in the natural environment, it may not come as a surprise that we did not detect any salinity-specific adaptive changes in the present study. Thus, we assume that the observed increase in total population size at 7PSU over time is mediated through phenotypic adaptation, which can for instance be achieved through a remodelling of the Sec protein export machinery (Ishii et al., 2015). As salinity did not seem to be a target for selection in the present study, we conclude that the delayed resistance evolution seen at 7 PSU and in the fluctuating treatment are driven, as the model predicts, by growth rate and its impact on phage production and the resulting fitness costs.

Reduction in growth is a common consequence of sub-optimal environmental conditions (Balakrishnan, De Silva, Hwa, & Cremer, 2021). The predicted net-effect of bacterial growth on resistance evolution and phage persistence in bacterial populations suggests that also changes in other environmental parameters, which are becoming more and more severe due to climate change, will have similar effects on phage bacteria interactions. The prolonged persistence of filamentous phages in bacterial populations can have additional consequences also for other organisms. That is because filamentous phages can carry deadly toxins, e.g., the cholera toxin which can turn bacteria into deadly diseases (Waldor & Mekalanos, 1996).

## Conclusion

Combining experimental evolution, genomics, and mathematical modelling we show that phage epidemics are prolonged in sub-optimal environmental conditions, which reduce bacterial growth rate. With decreasing growth rate, phage production declines and the costs for phage carriage relative to surface receptor mutation becomes progressively smaller which ultimately relaxes selection for SRM. Based on our model, which identifies bacterial growth rate as the driving factor behind these sub-population dynamics, we suggest that not only salinity but also other environmental parameters, which reduce bacterial growth rate, will reproduce these findings. In times of environmental change, we predict that altered interactions and co-evolutionary dynamics between phages and bacteria will also change higher order interactions, which are often mediated by phages (Wendling et al., 2017). This can further by intensified when fast changes in abiotic parameters, such as salinity, require recourse allocation towards respective coping mechanisms, i.e., osmoregulation in host organisms which impairs their immune systems and ultimately increases the impact of infections (Goehlich, Sartoris, Wagner, Wendling, & Roth, 2021).

## Supporting information

Supplementary Materia

## Acknowledgements

We thank Katja Trübenbach, Kim-Sara Wagner, Diana Gill, and Veronique Merten for their support in the laboratory. We thank Melanie Heinemann and Mechthild Bömeke for excellent support in the sequencing laboratory

## Author contributions

HG: experiment, data analysis and interpretation, first draft, final draft

OR: funding, study design, data interpretation, final draft

MS: mathematical model, contributed to first draft, final draft

CMC: genomic analysis, final draft

AP: genomic analysis, final draft

JR: genomic analysis and interpretation, final draft

HL: genomic analysis and interpretation, final draft

CCW: funding, study design, data analysis and interpretation, first draft, final draft

## Data deposit

All experimental data have been deposited on dryad: https://doi.pangaea.de/10.1594/PANGAEA.936135?format=html. Genomic data are available at NCB SubmissionID SUB12140318, and in the supplemental data file electronic supplementary material, Table S11.

