## Supplementary Materia for "SUB-OPTIMAL ENVIRONMENTAL CONDITIONS PROLONG PHAGE EPIDEMICS IN BACTERIAL POPULATIONS"

SUPPLEMENTARY MATERIAL

1 **Mathematical Model**

We employed the same model as presented in (Wendling et al., 2022). The dynamics of the ancestral, non-resistant clones (with density *B*), resistant SIE clones (*I*), resistant SRM clones (*R*), and SIE clones that have also acquired the MSHA mutation (*IR*), as well as the phage population (*V*) in batch cultures is modelled by the following system of differential equations:

$$\frac{dB}{dt}=\left( 1-m \right)g\left( B_{total} \right)B-\phi BV$$

$$\frac{dI}{dt}=\left( 1-m \right)\left( 1-v \right)g\left( B_{total} \right)I+\phi BV$$

$$\frac{dR}{dt}=\left( R+mB \right)g\left( B_{total} \right)$$

$$\frac{dIR}{dt}=\left( \left( 1-v \right)IR+mI \right)g\left( B_{total} \right)$$

$$\frac{dV}{dt}=ß\left( I+IR \right)-dV$$

Bacterial growth was modelled by generalized logistic growth of the form $g\left( B_{total} \right)=r\left( 1-\left( \frac{B_{total}}{K} \right)^{w} \right)$. Here *r* is the maximum growth rate (mgr) of the ancestral bacterial clone, *K* is the carrying capacity of the batch culture and $B_{total}=B+I+R+IR$ is the total density of all bacterial types. The curvature parameter *w* determines whether maximum growth is attained at an early point in the growth phase (*w* < 1) or at a late point (w > 1) . We assume that SIE clones (I and IR) suffer a growth rate reduction relative to the ancestral type due to virulence caused by intra-cellular production of virus, here represented by the virulence factor v <= 1. A completely avirulent virus would have v=0, and maximum virulence v=1 corresponds to growth arrest of the bacterial cell.

Viruses (*V*) infect non-resistant ancestral bacteria (*B)* following a mass action law with adsorption rate (phi), reflecting that increasing densities of either bacteria or viruses lead to higher encounter rates and thus more infections. Infection of a bacterial cell transforms cells into a resistant SIE clone (*I)*, which actively produces new viral particles (*V*) with phage production rate (*ß*), which we assume to be constant and independent of bacterial growth rate. Additionally, both ancestral bacteria (*B)* and SIE clones (*I)* can acquire complete resistance (*R* & *IR*) through mutations within the MSHA type IV pilus operon. We assume that MSHA-mutants have the same growth rate as the ancestor.

All bacterial types grow until the carrying capacity (*K*) is reached, but bacteria-virus interactions continue to occur as long as there are sensitive bacteria and viruses left. After a certain time t*max* a portion (here 1/100th) of the entire community is transferred to fresh medium and the process restarts.

2 **Tables**

**Table S1: Analysis of Variance Table (Type marginal – F-statistics) for overall generalised least square model (gls) of Cell forming units (CFU).** Degrees of freedom: 665 total; Residuals 653.

|  | **df** | **F-value** | **Pr(>F)** |
| --- | --- | --- | --- |
| Intercept | 1 | 22.0 | < 0.001 |
| Phage | 1 | 51.4 | < 0.001 |
| Salinity | 2 | 3.3 | 0.039 |
| Transfer | 1 | 83.2 | < 0.001 |
| Phage:Transfer | 1 | 36.7 | < 0.001 |
| Phage:Salinity | 2 | 3.0 | 0.048 |
| Salinity:Transfer | 2 | 1.0 | 0.089 |
| Phage:Salinity:Transfer | 2 | 5.2 | **0.006** |

**Table S2: Analysis of Variance Table (Type marginal – F-statistics) for simplified overall generalised least square model (gls) of viron dynamics in evolution experiment. Model simplification of model above according to AIC.** Degrees of freedom: 678 total; Residuals 670.

|  | **df** | **F-value** | **Pr(>F)** |
| --- | --- | --- | --- |
| Intercept | 1 | 78052.4 | < 0.001 |
| Phage | 1 | 64.0 | < 0.001 |
| Salinity | 2 | 8.5 | < 0.001 |
| Transfer | 1 | 43.2 | < 0.001 |
| Phage:Transfer | 1 | 39.7 | < 0.001 |
| Phage:Salinity | 2 | 3.3 | **0.039** |

**Table S3: Analysis of Deviance Table (Type II Wald chisquare tests) for reduced generalised mixed effect model (glmer) of resistance (Reduction in bacteria growth assay (RBG).** Testing Number of observations: 303. Binomial error distribution.

|  | **df** | **Chisq** | **p value** |
| --- | --- | --- | --- |
| Transfer | 1 | 62.8 | < 0.001 |
| Salinity | 2 | 6.1 | 0.047 |
| Phage | 1 | 176.6 | < 0.001 |
| Transfer:Salinity | 2 | 9.8 | **0.008** |
| Transfer:Phage | 1 | 8.1 | **0.004** |
| Salinity:Phage | 2 | 8.9 | **0.011** |

**Table S4** Parameter values of the mathematical model and their biological meaning

| **Parameter** | **Biological meaning** | **Value** |
| --- | --- | --- |
| *r* | Maximum growth rate (mgr) of ancestor *B* | 15 PSU: 3 (h^-1^)  7 PSU: 1.5 (h^-1^) |
| *K* | Carrying capacity of bacteria | 10^9^ cells/ml |
| *w* | Curvature parameter | 0.02 (Wendling et al., 2022) |
| *v* | Virulence | 0.3 |
| *phi* | Phage adsorption rate | 10^-8^ (h^-1^) (Taddei & De Paepe, 2006) |
| *ß* | Phage production rate | 60 (phages/cell h^-1^) |
| *d* | Phage decay rate | 0.01 (h^-1^) (Taddei & De Paepe, 2006) |
| *m* | Mutation rate | 10^-8^ |

**Table S5: Analysis of Variance Table (Type marginal – F-statistics) for generalised least square model (gls) of Cell forming units (CFU) using control populations only.** Degrees of freedom: 313 total; Residuals 307

|  | **df** | **F-value** | **Pr(>F)** |
| --- | --- | --- | --- |
| Intercept | 1 | 211.4 | < 0.001 |
| Salinity | 2 | 10.9 | < 0.001 |
| Transfer | 1 | 0.1 | 0.712 |
| Salinity:Transfer | 2 | 3.7 | **0.026** |

**Table S6: Analysis of Variance Table (Type marginal – F-statistics) for generalised least square model (gls) of Cell forming units (CFU) using co-evolving populations only.** Degrees of freedom 352 total; Residuals 346.

|  | **df** | **F-value** | **Pr(>F)** |
| --- | --- | --- | --- |
| Intercept | 1 | 11.7 | < 0.001 |
| Salinity | 2 | 1.7 | 0.193 |
| Transfer | 1 | 121.6 | < 0.001 |
| Salinity:Transfer | 2 | 3.6 | **0.030** |

**Table S7: Analysis of Variance Table (Type marginal – F-statistics) for generalised least square model (gls) of virons dynamics in evolution experiment.** Dataset includes only co-evolving populations from T3-T09. Degrees of freedom: 354 total; Residuals 348.

|  | **df** | **F-value** | **Pr(>F)** |
| --- | --- | --- | --- |
| Intercept | 1 | 1116222.7 | < 0.001 |
| Salinity | 2 | 8.7 | < 0.001 |
| Transfer | 1 | 1.1 | < 0.001 |
| Salinity:Transfer | 2 | 6.0 | **0.004** |

**Table S8: Analysis of Deviance Table (Type II Wald chisquare tests) for generalised mixed effect model (glmer) of resistance (Reduction in bacteria growth assay (RBG) using only co-evolving populations.** Number of observations: 172. Binomial error distribution.

|  | **df** | **Chisq** | **p value** |
| --- | --- | --- | --- |
| Transfer | 1 | 64.6 | < 0.001 |
| Salinity | 2 | 7.3 | 0.026 |
| Transfer:Salinity | 2 | 9.7 | **0.008** |

**Table S8a: Post-hoc Tukey Contrasts,** Resistance of bacteria in co-evolving populations

|  | **Estimate** | **Std. Error** | **t value** | **Pr(>\|t\|)** |
| --- | --- | --- | --- | --- |
| 15 PSU – 7 PSU == 0 | -0.37 | 0.10 | -3.9 | < 0.001 |
| Fluctuating – 7 PSU == 0 | -0.32 | 0.01 | -3.3 | 0.002 |
| Fluctuating – 15 PSU == 0 | 0.05 | 0.09 | 0.6 | 0.838 |

**Table S9: Analysis of Deviance Table (Type II Wald chisquare tests) for generalised mixed effect model (glmer) of resistance mode (PCR) using only infected populations.** Number of observations: 128. Binomial error distribution.

|  | **df** | **Chisq** | **p value** |
| --- | --- | --- | --- |
| Transfer | 1 | 489.9 | < 0.001 |
| Salinity | 2 | 80.4 | < 0.001 |
| Transfer:Salinity | 2 | 32.0 | **< 0.001** |

**Table S10: Post-hoc Tukey Contrasts, Resistance of bacteria in co-evolving populations**

|  | **Estimate** | **Std. Error** | **t value** | **Pr(>\|t\|)** |
| --- | --- | --- | --- | --- |
| 7 PSU – 15 PSU == 0 | -3.52 | 0.42 | -8.4 | < 0.001 |
| Fluctuating – 15 PSU == 0 | -1.22 | 0.31 | -4.0 | < 0.001 |
| Fluctuating – 7 PSU == 0 | 2.30 | 0.44 | 5.2 | < 0.001 |

**Table S11 SNV Table of all clones evolved in the presence of the filamentous phage VALGΦ8 at either 15 PSU, 7 PSU, or in the fluctuating treatment**. SNVs occurring within the borders of the mshA operon conferring phage resistance are indicated in bold. HP = hypothetical protein. Res = resistance mode, Sal = Salinity during evolution experiment. Φ_M: Φ-particle associate mutants

*1: N_acetylgalactosamine_N, diacetylbacillosaminyl_diphospho_undecaprenol_4_alpha_N_acetylgalactosaminyltrans-ferase/O_antigen_biosynthesis_glycosyltransferase_WbnH

*2: N_diacetylbacillosaminyl_diphospho_undecaprenol_4_alpha_N_acetylgalactosaminyl-transferase/O_antigen

_biosynthesis_glycosyltransferase_WbnH

| **Clone** | **Res** | **Sal** | **Position** | **Type** | **Description** | **Region** |
| --- | --- | --- | --- | --- | --- | --- |
| **VALG076** | **SRM** | **7** | **3011353** | **E1405*_(GAA_TAA)** | **HP** | **K01M1_28140** |
|  |  |  | 3238929 | noncoding_(13/1550nt) | 16S_ribosomal_RNA | K01M1_30280 |
| **VALG079** | **SRM** | **7** | **3011167** | **W1467G_(TGG_GGG)** | **HP** | **K01M1_28140** |
| **VALG084** | **SRM** | **15** | **3011082** | **coding_(4484/4527nt)** | **HP** | **K01M1_28140** |
| **VALG085** | **SRM** | **15** | **3021814** | **coding_(639_649/1725nt)** | **Type_II_secretion_system_protein_E** | **epsE_2** |
| **VALG088** | **SRM** | **15** | **3024873** | **coding_(1199_1208/1641nt)** | **Type_3_secretion_system_secretin** | **sctC_2** |
|  |  |  | 3238929 | noncoding_(13/1550nt) | 16S_ribosomal_RNA | K01M1_30280 |
| **VALG090** | **SRM** | **15** | **3012163** | **coding_(3403/4527nt)** | **HP** | **K01M1_28140** |
| VALG092 | SRM | 15 | 2924788 | T216M_(ACG_ATG) | Twitching_mobility_protein | pilT_2 |
|  |  |  | 3238929 | noncoding_(13/1550nt) | 16S_ribosomal_RNA | K01M1_30280 |
| **VALG094** | **SRM** | **15** | **3015870** | **coding_(177/477nt)** | **HP** | **K01M1_28150** |
| **VALG100** | **SRM** | **FL** | **3024862** | **S407P_(TCC_CCC)** | **Type_3_secretion_system_secretin** | **sctC_2** |
| **VALG071** | **SIE** | **7** | **3012432** | **coding_(3126_3134/4527nt)** | **HP** | **K01M1_28140** |
|  |  |  | 3238929 | noncoding_(13/1550nt) | 16S_ribosomal_RNA | K01M1_30280 |
| VALG072 | SIE | 7 | 1701158 | intergenic_(+348/+304) | Sensor_histidine_kinase_RcsC/HP | rcsC_3_/K01M1_16000 |
|  |  |  | 1701160 | intergenic_(+350/+302) |  |  |
|  |  |  | 1701356 | intergenic_(+546/+106) |  |  |
| VALG073 | SIE | 7 | 3238929 | noncoding_(13/1550nt) | 16S_ribosomal_RNA | K01M1_30280 |
| VALG077 | SIE | 7 | 1701158 | intergenic_(+348/+304) | Sensor_histidine_kinase_RcsC/HP | rcsC_3_/K01M1_16000 |
|  |  |  | 1701160 | intergenic_(+350/+302) |  |  |
|  |  |  | 2614712 | K276Q_(AAA_CAA) | HP | K01M1_24360 |
|  |  |  | 3238929 | noncoding_(13/1550nt) | 16S_ribosomal_RNA | K01M1_30280 |
| VALG080 | SIE | 7 | 223607 | intergenic_(+2/_17) | *1 | *2 |
|  |  |  | 2033390 | L263H_(CTT_CAT) | Virulence_sensor_protein_BvgS | bvgS |
|  |  |  | 3238929 | noncoding_(13/1550nt) | 16S_ribosomal_RNA | K01M1_30280 |
| VALG081 | SIE | 7 | 3238929 | noncoding_(13/1550nt) | 16S_ribosomal_RNA | K01M1_30280 |
| **VALG082** | **SIE** | **7** | **3011389** | **coding_(4171_4177/4527nt)** | **HP** | **K01M1_28140** |
| VALG086 | SIE | 15 | 1701158 | intergenic_(+348/+304) | Sensor_histidine_kinase_RcsC/HP | rcsC_3_/K01M1_16000 |
|  |  |  | 1701160 | intergenic_(+350/+302) |  |  |
|  |  |  | 1701355 | intergenic_(+545/+104) |  |  |
|  |  |  | 1701369 | intergenic_(+559/+93) |  |  |
|  |  |  | 1701374 | intergenic_(+564/+88) |  |  |
|  |  |  | 1701376 | intergenic_(+566/+86) |  |  |
|  |  |  | 1701384 | intergenic_(+574/+77) |  |  |
|  |  |  | 1701391 | intergenic_(+581/+70) |  |  |
|  |  |  | 1701397 | intergenic_(+587/+65) |  |  |
|  |  |  | 1701400 | intergenic_(+590/+62) |  |  |
|  |  |  | 1701407 | intergenic_(+597/+55) |  |  |
|  |  |  | 1701410 | intergenic_(+600/+52) |  |  |
|  |  |  | 1701413 | intergenic_(+603/+49) |  |  |
| **VALG091** | **SIE** | **15** | **3021309** | **P385Q_(CCA_CAA)** | **Type_II_secretion_system_protein_E** | **epsE_2** |
|  |  |  | 3238929 | noncoding_(13/1550nt) | 16S_ribosomal_RNA | K01M1_30280 |
| VALG093 | SIE | 15 | 1701158 | intergenic_(+348/+304) | Sensor_histidine_kinase_RcsC/HP | rcsC_3_/K01M1_16000 |
|  |  |  | 1701160 | intergenic_(+350/+302) |  |  |
| VALG095 | SIE | FL | 1701158 | intergenic_(+348/+304) | Sensor_histidine_kinase_RcsC/HP | rcsC_3_/K01M1_16000 |
|  |  |  | 1701160 | intergenic_(+350/+302) |  |  |
|  |  |  | 1701356 | intergenic_(+546/+106) |  |  |
|  |  |  | 1701358 | intergenic_(+548/+104) |  |  |
| VALG097 | SIE | FL | 27910 | intergenic_(+62/_263) | tRNA_Val/23S_ribosomal_RNA | K01M1_00270_/_K01M1_00280 |
|  |  |  | 28000 | intergenic_(+152/_173) |  |  |
|  |  |  | 28036 | intergenic_(+188/_137) |  |  |
|  |  |  | 1701158 | intergenic_(+348/+304) | Sensor_histidine_kinase_RcsC/HP | rcsC_3_/K01M1_16000 |
|  |  |  | 1701160 | intergenic_(+350/+302) |  |  |
|  |  |  | 1701356 | intergenic_(+546/+106) |  |  |
|  |  |  | 1701358 | intergenic_(+548/+104) |  |  |
|  |  |  | 2427429 | coding_(131/1053nt) | HP | K01M1_22510 |
| VALG101 | SIE | FL | 1701158 | intergenic_(+348/+304) | Sensor_histidine_kinase_RcsC/HP | rcsC_3_/K01M1_16000 |
|  |  |  | 1701160 | intergenic_(+350/+302) |  |  |
| **VALG096** | Φ**_SRM** | **FL** | **3024844** | **Q413*_(CAA_TAA)** | **Type_3_secretion_system_secretin** | **sctC_2** |
| **VALG098** | Φ**_SRM** | **FL** | **3014037** | **W510*_(TGG_TAG)** | **HP** | **K01M1_28140** |
| **VALG102** | Φ**_SRM** | **FL** | **3022619** | **W312*_(TGG_TAG)** | **HP** | **K01M1_28240** |
| **VALG103** | Φ**_SRM** | **FL** | **3012432** | **coding_(3126_3134/4527nt)** | **HP** | **K01M1_28140** |
| **VALG104** | Φ**_SRM** | **FL** | **3022521** | **Q345*_(CAA_TAA)** | **HP** | **K01M1_28240** |

3**Figures**

**Supplement Figure 1**: **24h growth curves in the absence (left) and presence of *Vibrio* phage VALGΦ8 averaged over all replicate populations per treatment** (n=6, ± s.e). Sub-samples of evolving populations were taken at Timepoint 1 of the evolution experiment.

**Supplement Figure 2: Population dynamics in the absence of VALGΦ8 over 30 transfers.** **(A)** Bacterial densities are shown as colony forming units (log_10_ CFU ml^-1^) and **(B)** Phage densities as phage particles (PP) measured by PEG precipitation (log_10_ PP ml^-1^) over 30 transfers (x-axis). The black, dashed, horizontal line in panel B indicates the quantification limit for phage concentrations. **(C)** Fraction of phage susceptible clones (n=24): Susceptibility was tested against the ancestral phage VALGΦ8. **(D)** Fraction of Φ-carrier within resistant clones. In all panels, individual replicate populations are represented by small dots (n=6) and means by larger points (± s.e.). Colors represent the salinity level during the evolution experiment (dark blue: 15 PSU, light blue: 7 PSU, blue and dashed line: fluctuating).

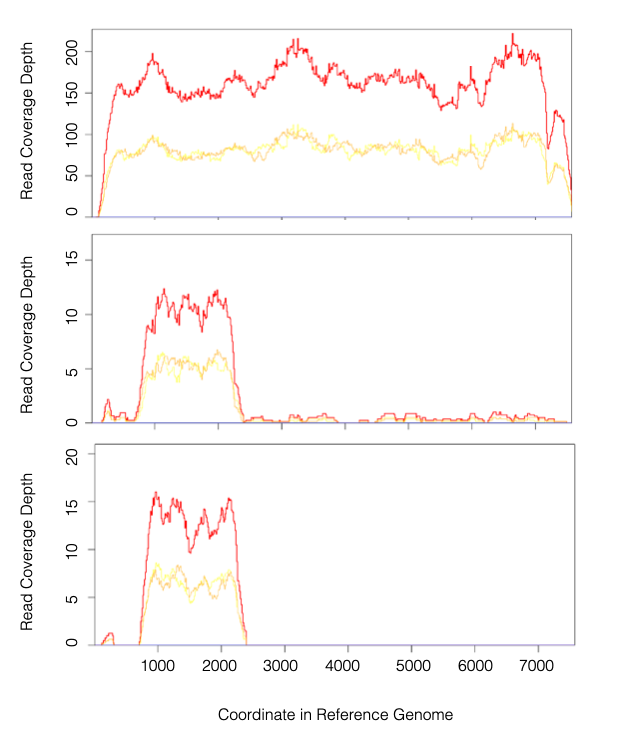

**Reads mapping to VALGΦ8**. Top: PCR-positive clones, reads map the entire genome of the infecting VALGΦ8. Bottom: PCR-negative Clones, here VALGΦ6 reads map to a shared region between the ancestral phage VALGΦ6 and the infecting phage VALGΦ8. Middle: PCR negative clones identified as Φ-particle associated mutants (Φ _SRM). The low number of reads that map over the complete VALGΦ8-genome indicates the presence of complete VALGΦ8-phage particles probably associated to the host cells.

**Figure S4: Theoretical population dynamics** modelled for 7 (light blue) and 15 PSU (dark blue), corresponding to growth rates of 1.5 1/h and 3 1/h, respectively. (a) bacterial densities in [CFU/ml], (b) phage densities in [PFU/ml]. (c) Fraction of phage susceptible population. Susceptible clones die out more quickly in 15 PSU. (d) Fraction of SIE clones within resistant subpopulation. Higher phage production resulted in faster spread of SRM and loss of SIE at 15 PSU.

4 **References:**

Taddei, F., & De Paepe, M. (2006). Viruses' life history: Towards a mechanistic basis of a trade-off between survival and reproduction among phages (vol 4, pg 1248, pg 2006). *PLOS Biology, 4*(8), 1470-1470. doi:ARTN e27310.1371/journal.pbio.0040273

Wendling, C. C., Lange, J., Liesegang, H., Sieber, M., Pohlein, A., Bunk, B., . . . Brockhurst, M. A. (2022). Higher phage virulence accelerates the evolution of host resistance. *Proc Biol Sci, 289*(1984), 20221070. doi:10.1098/rspb.2022.1070
